# Investigation of Codon Alternation Patterns in Genetic Diseases through Numerical Representation and Codon Classification

**DOI:** 10.1101/2020.03.02.971036

**Authors:** Antara Sengupta, Subhadip Chakraborty, Pabitra Pal Choudhury, Swarup Roy, Jayanta Kumar Das, Ditipriya Mallick, Siddhartha S Jana

## Abstract

Alteration of amino acids is possible due to mutation in codons that could have potential reasons to occur disease. Single nucleotide substitutions (SNS) in genetic codon thus have prime importance for their ability to occur mutations that may be deleterious indeed. Effective mutation analysis can help to predict the fate of the diseased individual which can be validated later by in-vitro experiments. Hence in this present study, we try to investigate the codon alteration patterns and their impact during mutation for the genes known to be responsible for a particular disease. We use a numerical representation of four nucleotides based on the number of hydrogen bonds in their chemical structures and make a classification of 64 codons as well as corresponding 20 amino acids into three different classes (Strong, Weak and Transitional). The entire analysis has been carried out based on these classifications. For our current study, we consider two neurodegenerative diseases, Parkinson’s disease, and Glaucoma. Several evidences claim similarities between both the diseases but proper pathogenetic factors are still unknown. The analysis reveals that the strong class of codons is highly mutated followed by the weak and transitional class. We observe that most of the mutations occur in the first or second positions in the codon rather than the third and mutations that occurred at the second place of codons are majorly deleterious. In most cases, the change in the determinative degree of codon due to mutation is directly proportional to the physical density property. Furthermore, we derive a determinative degree of five wild-type amino acid sequences, which can help biologists to understand the evolutionary relationship among them based on amino acid occurrence frequencies in proteins. In this regard we proposed an alignment-free method **SSADDA** (**S**equence **S**imilarity **A**nalysis using **D**eterminative **D**egree of **A**mino acid). Thus, our scheme gives a more microscopic and alternative representation of the existing codon table that helps in deciphering interesting codon alteration patterns during mutations in disease pathogenesis.

## 1. Introduction

Genes are the functional units of heredity [1]. It is mainly responsible for the structural and functional changes and for the variation in organisms which could be good or bad. Gene expressions are largely controlled by codon usage. Surprisingly it happens mainly due to the effects on transcription [2]. Hence, any mishaps during DNA transcription might lead to an adverse change in the genetic code which alters the protein synthesis. Single nucleotide substitutions (SNS) in genetic codon have prime importance for their ability to occur mutations [3]. Base substitutions can have a variety of effects [4]. The silent mutation is an example of a base substitution, where the change in nucleotide base has no outward effect, and are evolutionarily neutral [5]. But for missense mutation, a single nucleotide substitution in the genetic codon can be responsible for a mutation that is deleterious and remarkably decrease protein stability [6]. Thus, a mutated protein that differs from its wild type by only one amino acid (due to mutation) can cause the disease phenotype [7]. The alteration of amino acid due to the alternation in codon during disease may not be an arbitrary phenomenon [8]. They follow a certain pattern of an alternation. Diseases may have the specific signature of codon alteration. Hence, it is an interesting issue to investigate any hidden or priorly unknown alternation patterns (if any, during mutation) lead to the occurrence of a particular disease. This may help geneticists to understand the mechanism of mutation in disease phenotype better. Moreover, some research articles state that three codon positions define different roles and these are important features while studying standard genetic code (SGC) [9]. According to them, mutations in individual codon positions have different impacts [10, 11, 12]. Five synonymous substitutions are possible in the first codon position, whereas the third one is the most degenerated position. The second codon position is the most conserved position [9]. Hence, to elucidate the impact of codon alteration or pattern of alteration, the impact of three codon positions is also equally important. To make the quantitative and computational analysis easy, it is essential to make a suitable numerical representation of those biological facts, features, or characteristics.

A plethora of contributions are dedicated for characterizing genes through the light of numerical representations [13, 14, 15, 16]. People have tried in different ways to construct a mathematical model for similarity analysis of nucleotide sequences [17, 16, 18]. Rumer in his work has introduced the existence of a symmetry in the genetic code, which is also known as Rumer’s transformation or Amino-Keto transformation [15]. The transformation method applied to codon changes the degeneracy of the codon from its original one. Gonzalez et al have constructed mathematical models of the genetic code [19, 20, 21, 22]. According to them Rumer’s symmetry could have originated in an ancestral version of the genetic code [23]. Numeric representation of each type of multiplet signifies the number of codons mapping to a particular amino acid. However, the above numerical representation gives less importance to hidden abilities of nucleotides and their positional impact in codon to participate in the mutations.

Sequence homology and sequence similarity are not same. Sequence homology refers to a statement about common evolutionary ancestry of two sequences, while similarity refers to the degree of likeness between two sequences. Sequence similarity is a measure of a factual relationship between sequences. Sequence similarity analysis is one of the efficient ways to characterize genomic sequences and decipher different biological information embedded in them. The similarities can be analyzed by two ways, viz. alignment-based method and alignment-free method. Multiple alignments of related sequences sometimes may provide the most helpful information on its phylogeny, but it can produce incorrect results when applied to more divergent sequence rearrangements [24]. Some of the multiple alignment methods (MSA) align sequences based on the order in which they are received. The MSA methods give priority to closely related sequences to be aligned first. In the cases of less divergent sequences, by sharing a common ancestor they may be clustered separately [25] and they can be more accurately aligned, but provide incorrect phylogeny. Moreover, the time complexity of most of the methods is *O*(*n*^2^), where n refers to the length of the sequences. Alignment-free methods are natural frameworks to search for the patterns and properties embedded in biological sequences. The comparison can be defined as any method to make quantitative analysis of sequences and find similarity/dissimilarity among them that does not use alignment at any step of algorithm. Those are computationally less expensive as time complexity is O(n).

Parkinson’s disease is a progressive disease of the nervous system, occurs due to loss of dopaminergic neurons in the nigrostriatal pathway. It is the second most common neurodegenerative disorder after Alzheimar’s disease. PINK1, PARKIN and DJ1 are three main pathogenic genes [26, 27]. Glaucoma is a condition of increased ocular pressure within the eye, causing irreversible blindness [28]. It is a multifactorial disease, where genetic and environmental both factors are responsible for disease pathogenesis [29]. MYOC (myocilin) and CYP1B1 are two significant genes responsible for occurring this disease. Now-a-days it is categorised under neurodegenerative disease [30]. People having Glaucoma have greater risk to get Parkinson’s disease. Many reported research works claim similarities between Parkinson’s disease and Glaucoma. According to them oxidative stress and Mitochondrial dysfunction have significant roles in both the diseases [31, 32]. Over-expression of PARKIN protects retinal ganglion cells in glaucoma [33].

In this present work as the first step of our analysis, we introduced an alignment-free method **SSADDA** (**S**equence **S**imilarity **A**nalysis using **D**eterminative **D**egree of **A**mino acid). In this method we proposed a numerical scheme for representing codons in a more effective and meaningful manner, giving due importance to the nucleotides and their positions in a codon. At first, we calculated a determinative degree of each codon followed by classifying 64 codons into three different classes (weak, transitional, strong) based on the strength of nucleotides. We next calculated the degree of each amino acid based on the determinative degrees of constituent codons and classify 20 amino acids into the above classes. We proposed the determinative degrees of 64 codons followed by the degree of 20 amino acids and determinative degree of any arbitrary amino acid sequence. The overall idea of the numeric representation, classifications of codons and the amino acids are to determine the role of the nucleotides at each position of a codon during mutations causing fatal diseases and thus to understand the codon alteration pattern during the disease pathogenesis. We use our scheme to understand the pattern of codon alteration in two neurodegenerative diseases namely, Parkinson’s disease and Glaucoma [34] and their effects on the physicochemical properties of DNA primary sequential level as well as secondary structural level. The proximity between a set of any arbitrary amino acid sequences can be measured based on determinative degrees they have.

## 2. Methods and Materials

### 2.1. Methods

The physicochemical properties of the four nucleotides differ from each other. Hence, the strengths of nucleotides in DNA sequence level have a deep impact on protein formation and thus alteration may cause genetic diseases. It is therefore important to analyze the impact of positional alteration of nucleotides in codon during mutation. To specify their impact during codon formation it is worthy to have distinct quantification of each nucleotide based on some specific criteria [17]. In this present study, we use Rumer’s numerical representation of four nucleotides based on the number of hydrogen bonds in their chemical structures. Due to the degeneracy factor, more than one codon may code for a single amino acid, leading to the multiplet structure of amino acids [13]. Rumer [15] in his seminal work tried to explain the degeneracy with the help of first two constituent nucleotides of a codon, termed as *root* and classify the roots into three different classes, namely *Strong, Transitional* and *Weak*. Due to the multiplet structure of codons, codons coding for the same amino acid can be separated with regards to the roots. For example, if we consider *xyz* as a codon, then root *xy* can be separated from *z*. Thus, with four nucleotide bases, we get 4^2^ = 16 possible roots, which has been represented using a rhombic structure. According to Rumer four nucleotides (Adenine (A), Thymine (T), Guanine (G), Cytosine C)) have been arranged in decreasing chronological order according to their strength which is **C, G, T, A** [15, 35]. Rumer’s representation is good in analyzing the impact of the positional alteration. However, Rumer’s model is limited only to dual nucleotides. It is also incomplete in the sense that it fails to classify all the available codons and hereby not being able to extend its impact into DNA sequence level.

#### 2.1.1. Numerical representation of codon

We extend the existing dual nucleotide representation of nucleotides towards triplet or codon. We shift the concept from *root* to *root* + 1, where root holds the first two positions of a codon. The first two positions of a codon having greater importance to code any amino acid. Hence, due to mutation when a codon gets changed, there is a possibility of its root being shifted from one class to another. For example, CTA is a codon code for Leucine and it belongs to the strong class. Now if due to point mutation C at the first position is substituted by G, then the codon GTA which codes for Valine will be shifted to transitional class as shown in Figure 1.

**Figure 1:**
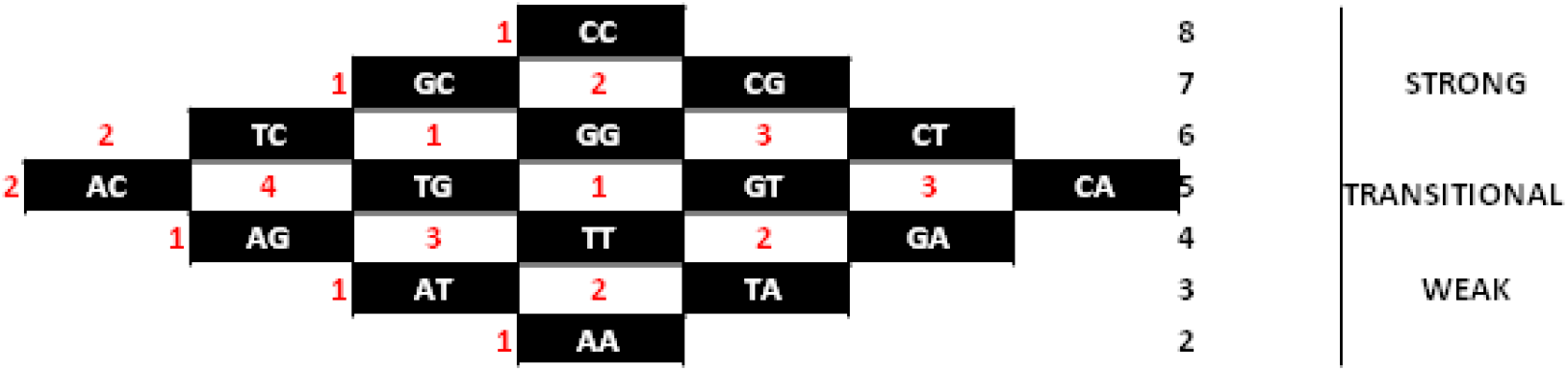
Rhombic structure of roots. Order of three dual nucleotide properties (Bottom to top): Weak, transitional and strong. The positional specifications of each dual nucleotide are recorded according to the average score of molecular weights of the amino acids they code).

We have represented each codon using a three digit number as shown in Figure 2. Each digit has its own significance. The physicochemical characteristics of the four nucleotides are not same. Mathematically DNA sequences are symbolic sequences. Hence, to make quantitative understanding of the impact of DNA during codon formation, it is necessary to specify their qualitative differences using quantitative value. In this present work we followed canonical ordering of C, G, U, A proposed by Rumer [15] in support with Duplij’s proposal to assign determinative degree to each nucleotide (an abstract characteristic of nucleotides) [35] where, *d*_*C*_ = 4, *d*_*G*_ = 3, *d*_*T*_ = 2, and *d*_*A*_ = 1. Thus, the first position of a codon indicates the additive degree of the root or class number, such that each dual nucleotide can have a range of numeric values from 2 to 8 only. For example, root **CC** scores 4 + 4 = 8; **AA** scores 1 + 1 = 2. If we consider Rumer’s rhombic arrangement of 16 dual nucleotides we can have seven groups of dual nucleotides (2, 3, 4, 5, 6, 7, 8) based on their additive scores. The second digit represents the position of the root in a particular row in the rhombic structure. We use positional specification at the second position of codon to provide a unique identity to each codon, else the additive degree of more than one codon would be the same. To represent a codon uniquely, we introduce a positional specification of each doublet or root in the rhombic organization of roots. To do so, we have calculated the average molecular weights of the corresponding amino acid(s) which starts with that particular dual nucleotide or root (shown in Table 1). The positions of dual nucleotides at each row are numbered accordingly. The last digit is the determinative degree of the third nucleotide. The format of numerical representation of codon table is shown in Figure 2.

**Table 1:** Positional Specification of root in the corresponding row in rhombic structure based on average molecular weight of the amino acids produced by them

| Dual Nucleotide | Amino Acid | Average Molecular Weight | Positional Specification in the Rhombic Structure |
| --- | --- | --- | --- |
| CC | P | 115.1 | 1 |
| GC | A | 89.1 | 1 |
| CG | R | 174.2 | 2 |
| GG | G | 75.1 | 1 |
| TC | S | 105.1 | 2 |
| CT | L | 131.2 | 3 |
| GT | V | 117.1 | 1 |
| AC | T | 139.65 | 2 |
| CA | H,Q | 150.7 | 3 |
| TG | C,W | 162.7 | 4 |
| AG | S,R | 139.65 | 1 |
| GA | D,E | 140.1 | 2 |
| TT | F,L | 148.2 | 3 |
| AT | I,M | 140.2 | 1 |
| TA | Y | 181.2 | 2 |
| AA | N,K | 139.15 | 1 |

**Figure 2:**
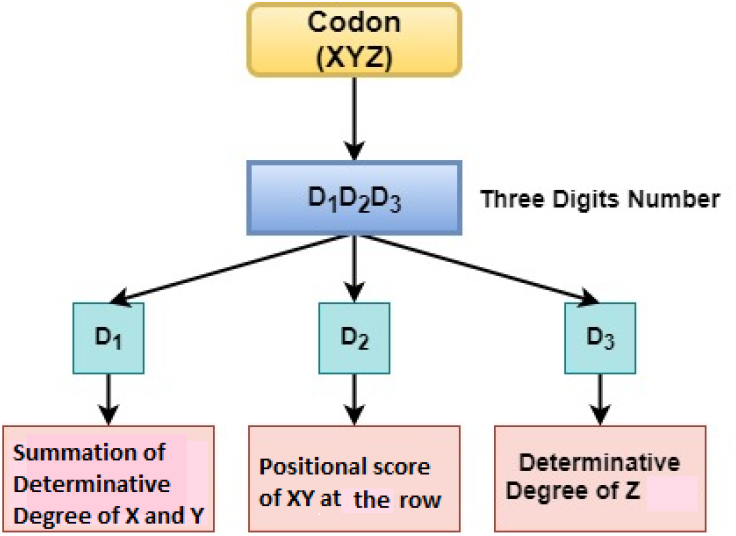
Scheme for three digits numerical representation of codons based on dual nucleotide scores and degenerative degree of third nucleotide.

As an example, for codon CCA, nucleotides C (having strength 4) at the first and second positions of the codon will form a dual nucleotide CC (4+4=8) at the first step. Thus the numeric value 8 is obtained at the first position of the determinative degree i.e. the root. The next step is to get the numeric value at the second position of the determinative degree, which is 1 because the root (CC) is placed at the first position of that particular row in the rhombic structure as specified in Figure 1. So now at the end of step 2, the value is 81. At the third step, it will select the numeric value 1 for the nucleotide at the third position, which is A. So now the determinative degree of the codon CCA is 811. It codes for the amino acid P (Proline). According to the classification of dual nucleotide, the CC comes under group G8 and the class of strong(S). We termed this three-digit representation of codon as the degree of codon and formally can be given as follows.

##### Definition 2.1

(Degree of Codon). *Let a codon C with constituent nucleotides say*, {*N*_1_, *N*_2_, *N*_3_}, *where N*_*i*_ ∈ *N, then the degree of C (δ*(*C*)*) is the concatenation (.) of three digits*.

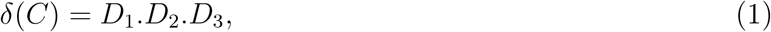

where, *D*_1_= determinative degree of dual nucleotide or additive degree of nucleotides, *D*_2_= the positional score of the root at particular row of the rhombic structure, and *D*_3_ is the determinative degree of third nucleotides as discussed above. Thus we derive determinative degrees of all 64 codons. The Table 2 (a) shows the standard genetic code table and the determinative degrees of all 64 codons are stated in Table 2(b).

**Table 2:**
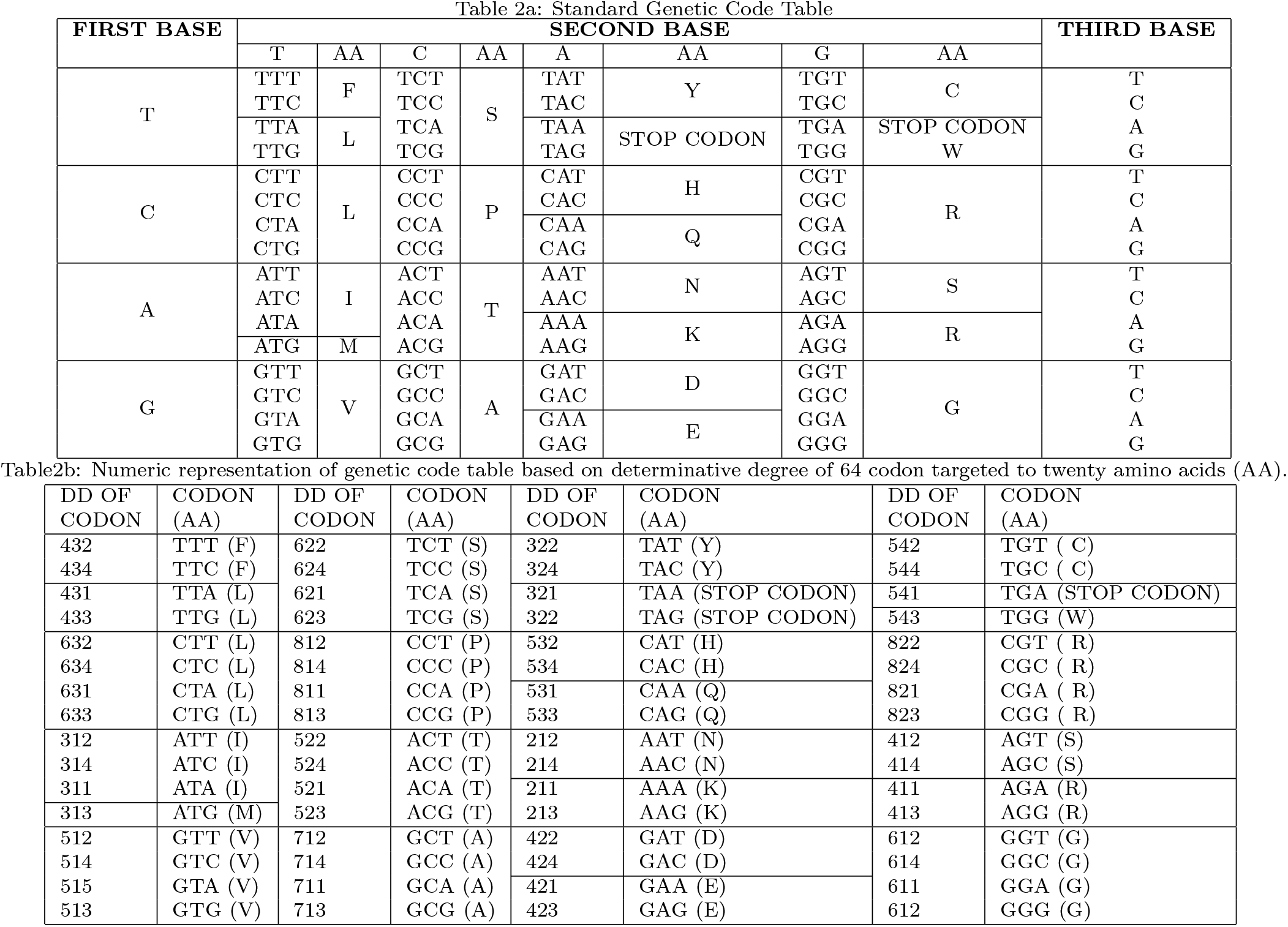
Determinative degree of 64 codons

The four nucleotides possess different bonding strengths based on the number of hydrogen bonds in their chemical structures. According to that **C** and **G** can be considered as strong bases whereas rest two i.e. **T** and **A** are weak bases. Again, within strong and weak base pairs they may have comparative strength within them such that pyrimidine bases, **C** and **T** are stronger than purine bases, **G** and **A**, respectively [36]. We classify all 64 codons into three classes namely Weak, Strong and Transitional according to the additive score of the root as shown in Figure 1. The 20 amino acids are also classified accordingly. We find six amino acids (P, A, R, S, G, L) as strong, five amino acids (T, C, W, H, Q) as transitional, and rest (F, L, D, E, I, M, Y, N, K, S, R) as weak. The maximum number of amino acids i.e. eleven (11) are fall under the weak class. Unlike, previous attempts we can classify all the codons into three different classes. While classifying, we observe an interesting fact that Arginine (R), Serine (S), and Leucine (L), each belong to two different classes, Weak and Strong. All three amino acids are coded by six (06) different codons, out of which four fall in the strong group and the rest of two in the weak group. The overlapping distribution of all the amino acids across three different classes is shown with the help of Ven-diagram in Figure 3.

**Figure 3:**
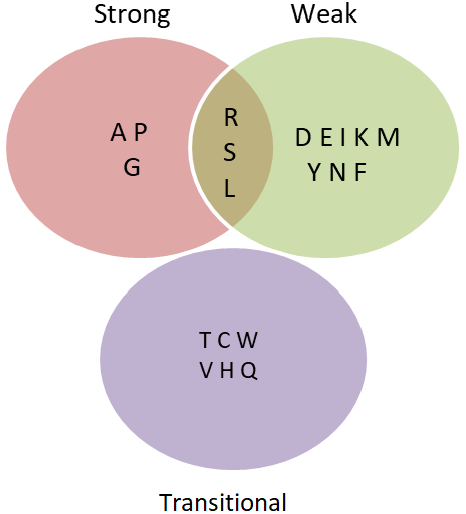
Distribution of 20 amino acids in three different overlapping classes.

#### 2.1.2. Classification of amino acids

#### 2.1.3. Determinative degree of amino acid

Due to degeneracy it is not possible to make one-to-one mapping between 64 codons and 20 amino acids, more than one codons are coding an amino acid. To handle the issue we calculate average determinative degree of all the constituent codons for an amino acid, termed as determinative degree of that amino acid. Let 𝒜 = {𝒜_1_, 𝒜_2_, · · ·, 𝒜_*n*_}be the set of twenty (*n* = 20) amino acids. To understand the impact of codons targeted to the amino acid, we calculate the degree (or strength) of amino acid by considering the average degree of all codons that code for a target amino acid.

##### Definition 2.2

(Degree of Amino Acid). *Let* 𝒜_*j*_ *be an amino acid where* 𝒜_*j*_ ∈ 𝒜 *and* 𝒜_*j*_ = {*C*_1_, *C*_2_, *C*_*n*_} *such that C*_*i*_ *is the constituent codon codes for* 𝒜_*k*_. *The degree of amino acid* 𝒜_*j*_ *denoted by δ’* (𝒜_*j*_) *can be calculated using the following equation:*

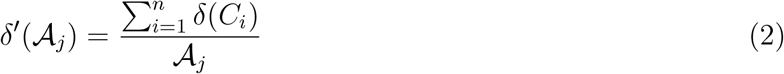

We report degree of all 20 amino acids calculated using Equation 2 in Table 3. Interestingly, it can be observed that the amino acids which come under strong class having determinative degree (DD) between the range of 612.5 to 812.5. In this way the range of DD for transitional class of amino acids lies between 512.5 to 543 and for weak class, it lies between 212 to 433. It is observed that Ser, Arg and Leu these three amino acids individually have two determinative degrees coming from two separate classes.

**Table 3:** Degree of twenty amino acids in three groups: Strong, Transitional and Weak.

| Amino Acid ($A_i$) | Class | $\delta'_{A_i}$ | Amino Acid ($A_i$) | Class | $\delta'_{A_i}$ |
| --- | --- | --- | --- | --- | --- |
| Lys | Weak Root | 212.00 | Ser | Weak Root | 413.00 |
| Asn |  | 213.00 | Glu |  | 422.00 |
| Tyr |  | 313.00 | Asp |  | 423.00 |
| Ile |  | 322.33 | Leu |  | 432.00 |
| Met |  | 323.00 | Phe |  | 433.00 |
| Arg |  | 412.00 |  |  |  |
| Val | Transitional Root | 513.50 | Leu | Strong Root | 612.50 |
| Thr |  | 522.50 | Gly |  | 632.25 |
| Gln |  | 532.00 | Ser |  | 622.50 |
| His |  | 533.00 | Ala |  | 712.50 |
| Cys |  | 543.00 | Arg |  | 822.50 |
| Trp |  | 543.00 | Pro |  | 812.50 |

#### 2.1.4. Determinative degree of amino acid sequence

Information about a protein’s function and its evolutionary history can be obtained from it’s primary structure or amino acid sequence. So it is essential to determine the protein’s amino acid sequence [37].

The frequency of occurrence of amino acids constituting an amino acid sequence varies from gene to gene and is well-conserved from species to species [38, 39]. The composition of protein works in such a way that the impact of mutations on protein structure can be minimized. Thus, amino acid composition of proteins reduces deleterious impact of mutations [40]. Moreover, amino acids have different propensities for forming alpha helices, beta sheets, and beta turns [37]. Hence, to understand the impact of occurrence of amino acids in an amino acid sequence on evolution, it is necessary to calculate the degree (or strength) of amino acid sequence by considering the average degree of all amino acids that constituents the amino acid sequence.

##### Definition 2.3

(Degree of amino acid sequence). *Let* 𝒮 = *A*_1_, *A*_2_, …*A*_*m*_ *be an arbitrary amino acid sequence of length m, where* 𝒜_*j*_ *be any amino acid, i.e. A*_*j*_ ∈ 𝒜. *The degree of amino acid sequence is denoted by δ’*(𝒮) *and can be calculated as follows:*

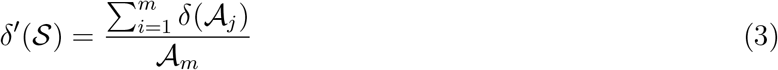

### 2.2. Collection of genome sequences

We consider five genes to carry out the experimentation. PARKIN, PINK1, DJ1 for Parkinson’s disease and, CYP1B1 and MYOC for Glaucoma. The wild type sequences of genes taken for Parkinson’s disease PARKIN (Accession No: AB009973.1), PINK1(Accession No: AB053323.1), DJ1 (Accession No: NM_007262.4), CYP1B1 (Accession No: NM_000104.3), MYOC (Accession No: NM_000261.1) are collected from NCBI GeneBank. We took a group of mutations reported in these genetic diseases collected from various online sources^§^,^§^,^§^, ^§^, ^§^. The Table 4 shows the summary of the mutated gene sequences collected.

**Table 4:**
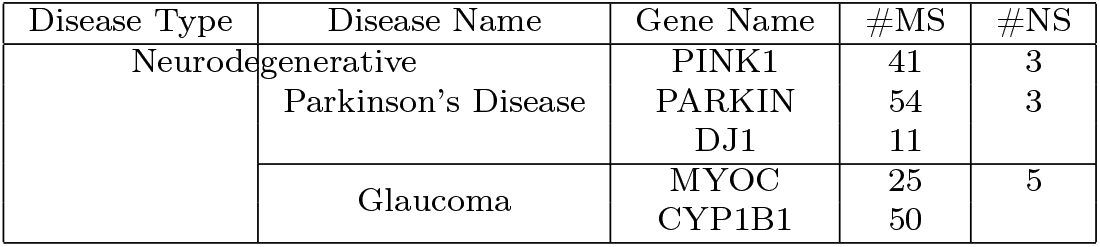
Mutation datasets (Missense sample (MS) and nonsense sample (NS)) of neuro-degenerative disease types and seven genes used for experimentation.

We consider only the point mutations (missense mutations, nonsense mutations). These are selected in an unbiased way based on the published reports for each mutation. More number the mutation reported, more strong the mutation to cause the disease. We considered at least three published reports for each mutation. However, in case of PINK1 and DJ1, we found very less reported evidences of mutation.

### 2.3. Time complexity analysis of alignment-free method SSADDA

The SSADDA method analyzes sequence similarity based on determinative degrees of their constituent amino acids. It is performed in two stages. In first stage, each triplet of the query sequence is read in terms of its determinative degree. Thus, for a sequence of length l it takes O(l) time to read and for n numbers of sequences it takes O(n×*l*) time. In second stage, each pair of amino acid sequences are compared based on the distribution of 20 amino acids in them and distance matrix is constructed. The time taken for this comparison is O(n^2^× 20). Hence, the total time taken to perform sequence similarity analysis is O(n×*l*)+O(n^2^ ×20) or O(nl)+On^2^. The time complexity of other contemporary alignment based methods like MAFFT is O(n^2^l+ nl^2^) [41], in Clustal Omega is O(n log n) [42], in Clustal W it is O(n^2^l^2^) [43], and in MUSCLE it is O(n^2^l) [43]. Hence, SSADDA is computationally more faster than other contemporary methods.

## 3. Results

Our prime objective is to analyze codon alteration patterns in disease phenotype. We consider two different types of neurodegenerative diseases for analysis, viz. Parkinson’s disease and Glaucoma. To make the computational analysis easy we represent codons numerically. With the help of our scheme, we analyze few mutated DNA sequences and compare their alteration pattern in comparison to non-diseased phenotypes. The experiments are therefore carried out in two ways, viz. quantitative characterization of mutations occurred and investigate evolutionary relationships among the wild type genome sequences of the genes responsible for the particular disease. For quantitative characterization of mutations occurred we perform few more experiments on the mutational data to understand various interesting facts and patterns that usually follow during mutation.

### 3.1. Codon class specific quantification (%) of mutation

Here in this subsection, it is aimed to investigate the general tendency of occurrence of mutation, whether the majority of mutations take place in codons of strong class or weak class or transitional class. The result has been shown in Figure 4. It can be observed from the figure that mutated samples of both types of diseases have a common tendency of having mutations in the strong class of codons, i.e. in G8, G7, or G6. In the case of PINK1, after strong class of codons (60.98%), the mutations have been taken place in the transitional class(24.39%). In the remaining four genes along with the strong class of codons, the weak class of codons have remarkable participation. In addition to that, we also checked the position of mutations in the context of a protein domain (active or nonactive) that may be altedred due to mutations. The active domains list is reported in Supplementary Table S1. Interestingly, we observe that mutations mostly present in the functional/active region. The overall observation also depicts that the strong class of codon is affected the most (Figure 4). We look into three amino acids Arginine (R), Serine (S), and Leucine (L), where each of them is coded by six codons, where four of six codons are from strong class and the remaining two codons are from weak class. We observe that the maximum mutated codons belong to the strong class except of gene PARKIN. In PARKIN mutated codons belong to amino acid ‘Serine’, which are from weak class. While checking the position of the mutations, we observe that mutations are from the active site domain of the protein, which may be altered due to mutations and ultimately lead to have pathogenic phenotype.

**Figure 4:**
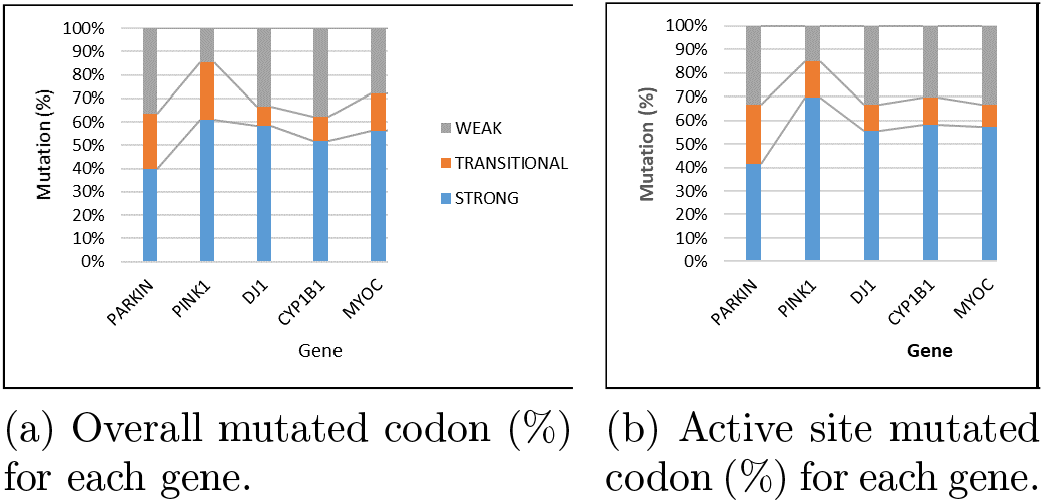
Gene wise percentage of mutated codons present in each class (Strong, Transitional and Weak) affected due to mutations.

### 3.2. Positional quantification of mutations in codons

Single point mutation may occur in any random position of codons. In a codon, out of three nucleotides, the first two nucleotides (from 5 end) usually determine the type of codon [44]. The role of third nucleotide appears to be very negligible. Our study with the mutated dataset of five genes once again proves the same fact (shows in Figure 5). Percentage wise calculations of mutations occurred at three different positions of codon state the tendency of mutations taking place at 1st and 2nd positions of codons. Comparatively, mutations at third nucleotides are very less with an exception in case of DJ1. In PARKIN, maximum mutations occurred at 1*st* position of the codons. In PINK1, CYP1B1 and MYOC, the mutations majorly occurred at the second position of the codon. The observation itself depicts the fact that the importance of the first two positions of a codon is more than third position during mutations.

**Figure 5:**
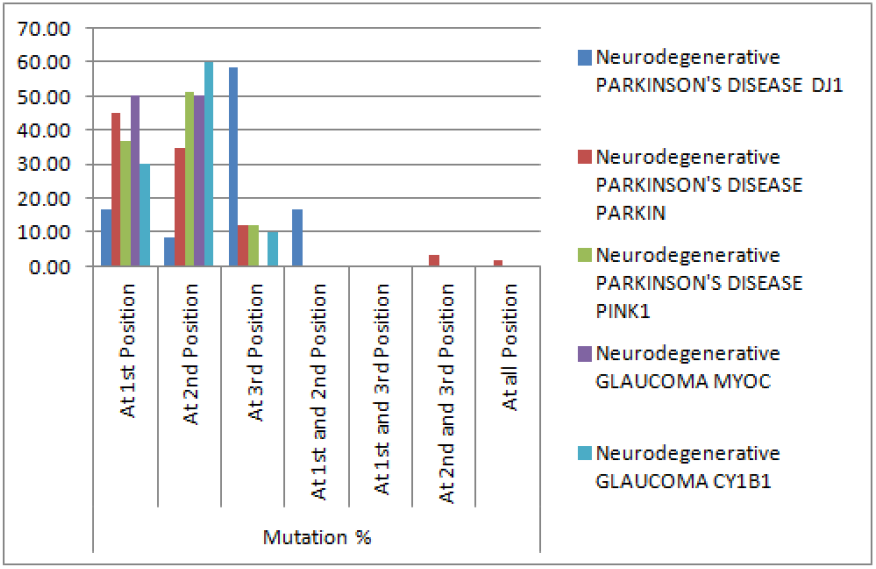
Mutations occurred (%) at different positions of codons.

### 3.3. Transition of codon groups due to mutation

Although mutations have majorly taken place at the codons belong to strong class, but there may be a possibility that during mutations the affected codon may transit from one group to another or may remain unchanged. Here in this subsection we try to find the pattern of group-wise transitions (G2 to G8) of codons occurred due to missense mutations. The transitions (in percentage) for all five genes are stated using directed graphs (shown in Figure 6).

**Figure 6:**
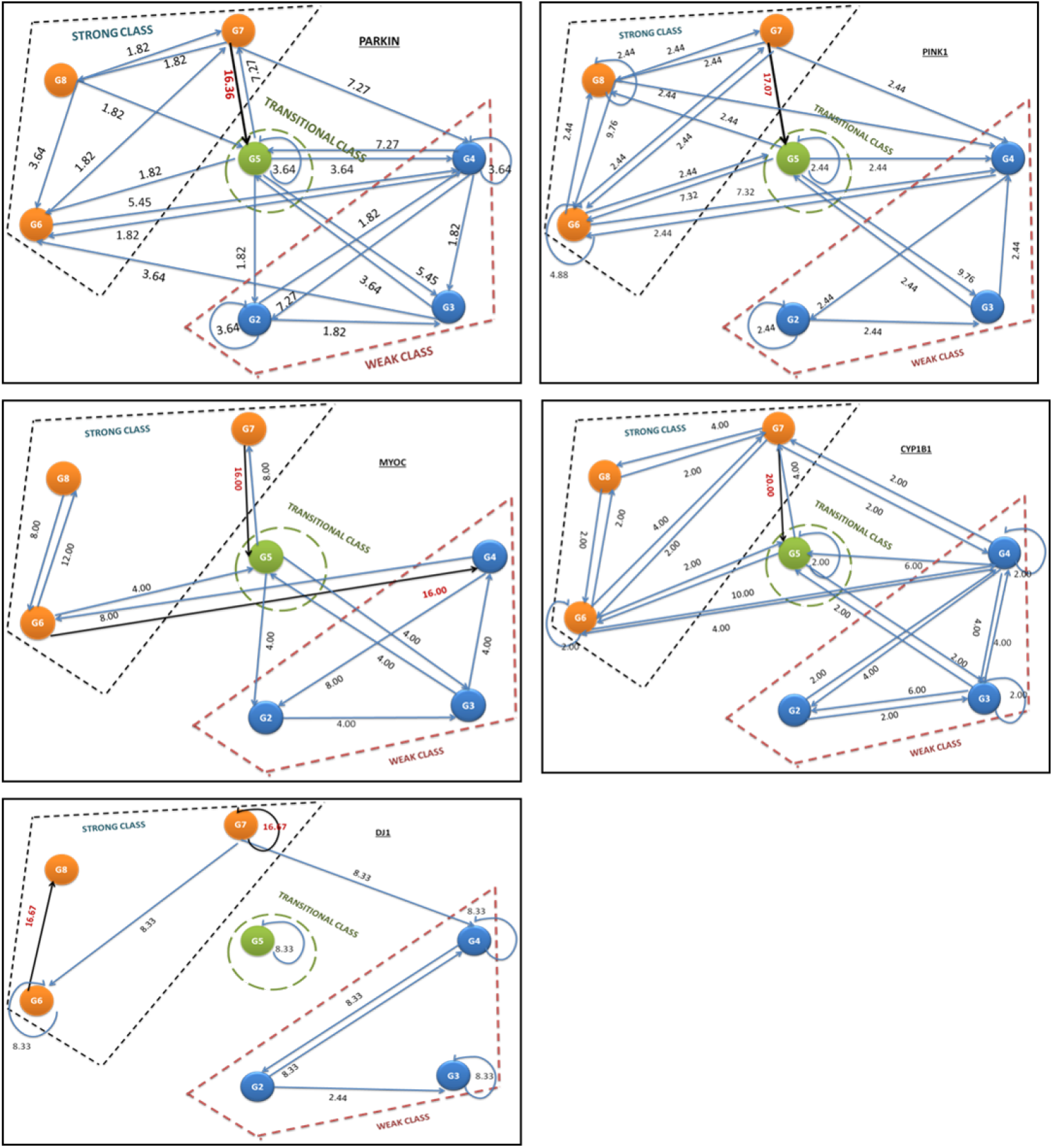
Transition of codon groups during mutations. The highest transitions are highlighted using red colour.

It can be observed that due to the missense mutations in genes responsible for neurodegenerative diseases (both Parkinson’s and Glaucoma), majority of mutations occurred at codons of group G7 (except DJ1, where we observe both in G7 and G6). We further observe that in most of the cases transitions due to mutations have taken places from G7 group to G5 group, except DJ1 where G7 group remains unaltered Supplementary Table S2. It implies that group G7 plays a significant role during mutations in neurodegenerative diseases. G7 contains the amino acids Ala and Arg, whereas, G5 contains His, Val, Trp, Glu, Cys, Thr.

### 3.4. Alteration of physical density of codon with the change in codon group due to mutation

Due to mutation, the physicochemical property like density of the whole gene sequence may get changes. To make a quantitative understanding of it, the density of each codon is deducted from the density of nucleotides, where the density of *C* = 1.55 *g/cm*^3^, *G* = 2.2 *g/cm*^3^, *T* = 1.32 *g/cm*^3^ and *A* = 1.60 *g/cm*^3^. Hence, the density of a codon is the average densities of the constituent nucleotides of the codon. We calculate the change (%) of density as shown in Figure 7. It is observed that in most of the cases density of codons due to mutations increased along with the transitions from weaker groups to stronger groups and decreased along with the transitions from stronger groups to weaker groups respectively (Supplementary Table S3).

**Figure 7:**
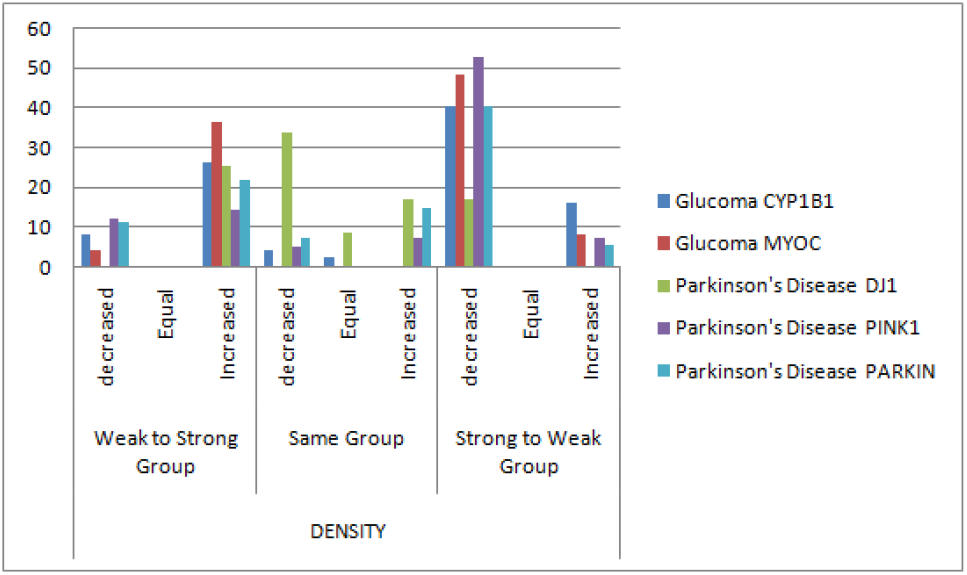
Trend of alteration in density of codon with the change in codon group due to mutation (%)

### 3.5. Investigating proximity between amino acid sequences based on their determinative degrees

Primary open-angle glaucoma (POAG) is the most common type of glaucoma. Reported research works hypothesized that POAG patients have a higher risk of getting Parkinson’s disease (PD) too [45]. Glaucoma is one of the common disease occurs of PD patients [46] and shows positive relationships between the disorders [47]. Hence in this subsection, it is tried to investigate proximity between five genes (namely, PARKIN, PINK1, DJ1, CYP1B1, MYOC) responsible for neurodegenerative diseases based on their determinative degree. To do so a distance matrix has been constructed to calculate the deviations between the determinative degrees of their wild-type amino acid sequences. Let determinative degree of any two arbitrary amino acid sequences *S*_1_ and *S*_2_ are *δ’*(*S*_1_) and *δ’*(*S*_2_), then the proximity between them with respect to their determinative degree is shown below:

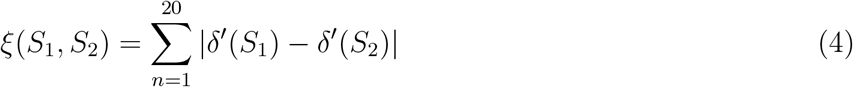

The distance matrix shown in Table 5 has been derived using equation (4) to measure proximity among the candidate amino acids. A phylogenetic tree is constructed from the distance matrix (shown in Figure 8. Thus, although PARKIN, PINK1, and DJ1 genes are responsible for Parkinson’s disease and CYP1B1 and MYOC are for Glaucoma, but the pictorial representation of the matrix shows a proximity between DJ1 and MYOC. Interestingly, it is also observed that genes MYOC and CYP1B1, although responsible for the same disease (Glaucoma) are distantly related in the phylogenetic tree.

**Table 5:** Proximity between amino acid sequences

|  | PARKIN | PINK1 | DJ1 | CYP1B1 | MYOC |
| --- | --- | --- | --- | --- | --- |
| PARKIN | 0.00 | 0.78 | 15.21 | 7.08 | 18.16 |
| PINK1 | 0.78 | 0.00 | 14.43 | 6.30 | 17.37 |
| DJ1 | 15.21 | 14.43 | 0.00 | 8.13 | 2.95 |
| CYP1B1 | 7.08 | 6.30 | 8.13 | 0.00 | 11.07 |
| MYOC | 18.16 | 17.37 | 2.95 | 11.07 | 0.00 |

**Figure 8:**
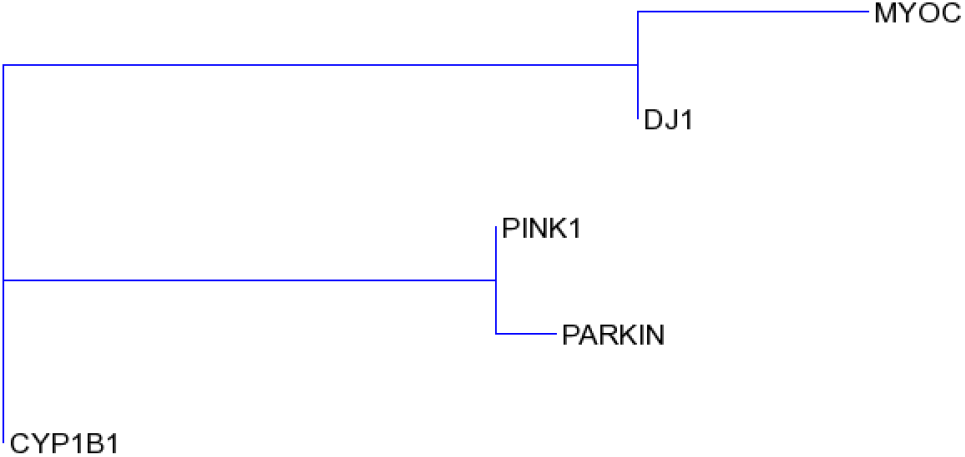
Phylogenetic tree showing evolutionary relationships among all five genes.

We compared our SSADDA method produced with those contemporary alignment-based methods based on the distance matrix derived for each of them. Pearson correlation coefficient (r) [48] is used to find correlations between each pair of matrices for comparison (as shown in Table 6). It has been found that correlation coefficient existing between few alignment-based methods Clustal Omega, MUSCLE, MAFFT) and SSADDA are more superior than that exist between ClustalW2 and other two alignment-based methods MUSCLE and MAFFT.

**Table 6:** Correlation coefficient between the matrices derived by four aligners (ClustalW, Clustal, MAFFT, MUSCLE) and SSADDA

|  | SSADDA | ClustalW2 | ClustalOmega | MAFFT | MUSCLE |
| --- | --- | --- | --- | --- | --- |
| **SSADDA** | **1** | 0.45 | 0.74 | 0.60 | 0.68 |
| **ClustalW2** |  | **1** | 0.65 | 0.45 | 0.62 |
| **Clustal Omega** |  |  | **1** | 0.92 | 0.98 |
| **MAFFT** |  |  |  | **1** | 0.94 |
| **MUSCLE** |  |  |  |  | **1** |

**Table 7:** Calculation of Robinson and Foulds Distance between each pair of phylogenetic trees

|  | SSADDA | CLUSTAL W2 | CLUSTAL Omega | MAFFT | MUSCLE |
| --- | --- | --- | --- | --- | --- |
| SSADDA | 0 | 4 | 2 | 2 | 4 |
| ClustalW2 |  | 0 | 4 | 4 | 2 |
| Clustal Omega |  |  | 0 | 4 | 4 |
| MAFFT |  |  |  | 0 | 2 |
| MUSCLE |  |  |  |  | 0 |

Phylogenetic trees are also constructed using other contemporary alignment-based methods viz. ClustalW2, Clustal Omega, MAFFT, and MUSCLE (shown in Figure 9). To make comparisons between the contemporary alignment-based methods and the alignment-free method SSADDA, Robinson and Foulds (RF) distance between each pair of phylogenetic trees are calculated (as shown in Figure 7), which shows more or less similar kind of results.

**Figure 9:**
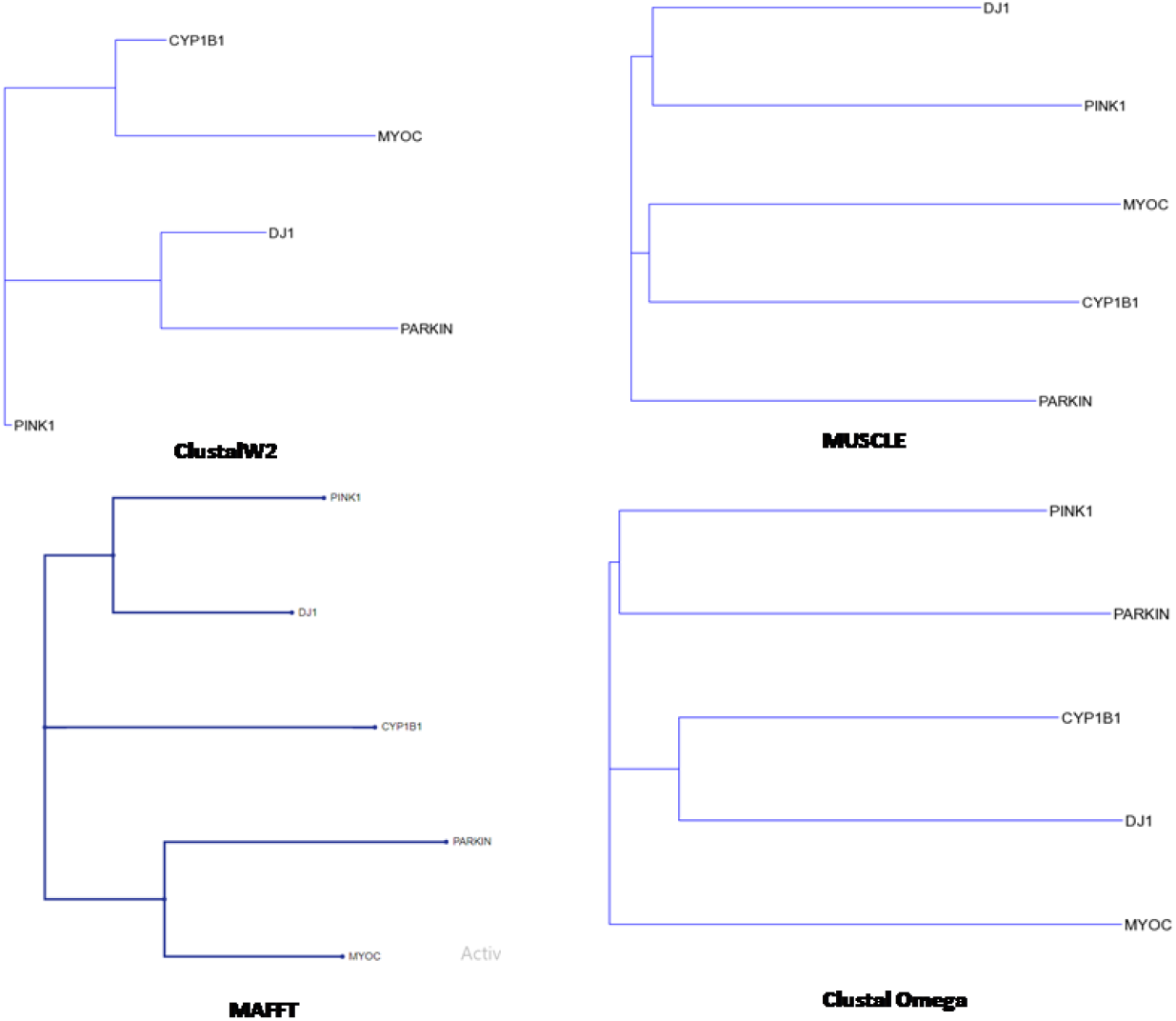
Phylogenetic trees are also constructed using different contemporary alignment-based methods.

### 3.6. Predicting the functional effect of mutations occurred

Mutations may be amino acid substitution or indel, the intensity of harmfulness may not be the same always. The mutation may or may not have an impact on the biological functions of a protein. PROVEAN (Protein Variation Effect Analyzer) is online software that predicts the functional effect of amino acid substitution [49]. The objective is to understand the effect of mutations in codon on the stability of the protein produced [50]. It is to be noted that isolates with a score equal to or below −2.5 are considered deleterious and scores above −2.5 are neutral. In this subsection gene-wise percentage of the deleterious mutations occurred are calculated using PROVEAN software (shown in Figure 10(a)). The graph shows maximum deleterious mutations taken place in MYOC and the chronological order is MYOC *>* CYP1B1 *>* PARK1 *>* PINK1 *>* DJ1, where MYOC and CYP1B1 both genes are responsible for Glaucoma. Figure 10 (b) shows the percentage of deleterious mutations that occurred due to the substitution of nucleotides at three different places of a codon. It is remarkable that in all the five genes deleterious mutations occur due to nucleotide substitution at the 2nd position of the codon. Some of the mutations reported in MYOC (V251A, P370L, S341P, G367R) are responsible for juvenile-onset open-angle glaucoma (JOAG), which affects people having age between 3 years to 40 years [51, 52, 53, 54]. According to our observations, they all are deleterious. Some of the deleterious mutations that occurred in CYP1B1 (R390H, G61E, Y81N, Y229K) are responsible for primary congenital glaucoma (PCG) [55, 56]. (D280N, R275W, R256C) are deleterious heterozygous parkin mutations take place in exon 7 and cause juvenile parkinsonism [57]. Some deleterious mutations that occurred in PINK1 like A168P, L347P, G409V, E240K interfere with auto-phosphorylation [58]. L10P and L166P both of the deleterious mutations remarkably decrease the stability of DJ1 protein [6]. Investigations have also been carried out to find the codon group transitions for deleterious mutations. It is observed that the deleterious mutations have majorly occurred in a strong group of codons. In mutations that occurred in the genes of PINK1, DJ1, and CYP1B1 the codon class remains unchanged, and in PARKIN and MYOC due to mutations, the codons have been changed from strong class to intermediate group (Table 8). Gene-wise deleterious mutations are shown in Supplementary Table S4.

**Table 8:**
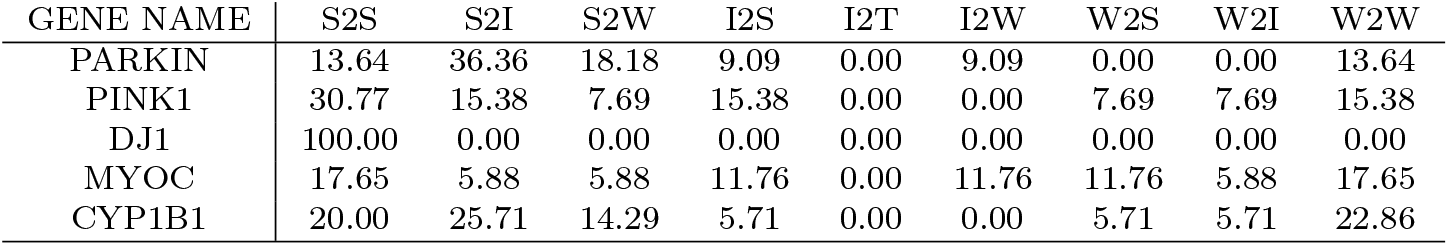
Transition of codon groups due to deleterious mutation in percentage

**Figure 10:**
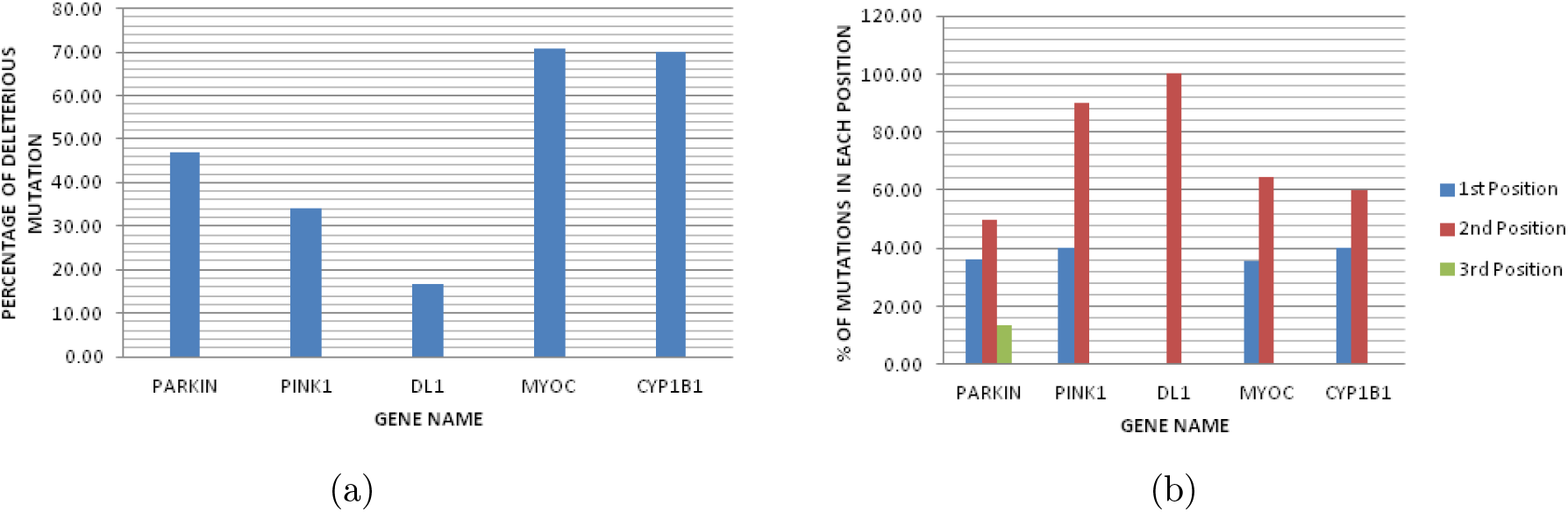
(a) Percentage of deleterious mutations occurred in all the genes. (b) Trend of alteration in different positions of codon(%) causing deleterious mutations

## 4. Discussion

In this present study, we try to investigate the codon alteration patterns and their impact during mutation for the genes known to be responsible for a particular disease. The physicochemical properties of the four nucleotides differ from each other. Hence, to specify their impact during codon formation it is worthy to have distinct quantification of each nucleotide based on some specific criteria [17]. We use Rumer’s numerical representation of four nucleotides based on the number of hydrogen bonds in their chemical structures and make a classification of 64 codons as well as corresponding 20 amino acids into three different classes (Strong, Weak and Transitional). According to the classification rule, the codons starting with (CC, CG, GC, TC, CT, GG) are from strong class, the codons starting with (AC, CA, TG, GT) are from transitional class and remaining (AG, GA, TT, AT, TA, AA) are from weak class. The genetic code table possesses three amino acids Arg, Ser, and Leu, where six codons specify a single amino acid. As a result, each of them poses two determinative degrees; One comes from a codon family (4 codons) and another from codon pair (2 codons). According to our mathematical scheme they belong to two different classes respectively. While the existing genetic code table only speaks about the name of the mutated amino acids, our rule of classification can give a more microscopic view. Using the classification rule we studied codon alteration patterns of missense mutations taken place in two different types of neurodegenerative diseases, namely Parkinson’s disease and Glaucoma. The analysis reveals that the strong class of codons are highly mutated followed by the weak and transitional class. The strong class of codons belong to groups G8, G7 and G6. We observe that most of the mutations occur in the first or second positions of the codons rather than the third and mutations that occurred at the second positions are of majorly deleterious. Moreover, the trend of alteration in density of codon with the change in codon group due to mutation shows chronological order *G*2 *< G*3 *< G*4 *< G*5 *< G*6 *< G*7 *< G*8. The analysis depicts that the changes in density of codons made due to mutations are directly proportional to the strength of codons. The similarities in experimental results indicate commonalities in selection of codons from respective genes for alteration, which cause the disease phenotype. Determinative degrees of five wild-type amino acid sequences (PARKIN, PINK1, DJ1, CYP1B1, MYOC) are derived using the determinative degrees of 20 amino acids. These distinct quantification of each amino acid sequence defines their passive characteristics based on the frequency of occurrence of amino acids in terms of their determinative degrees. The motivation here is to investigate the proximity between the amino acid sequences. The phylogenetic tree constructed shows proximity between DJ1 and MYOC, which again gives a holistic view of the interrelationships existing between the genes participating in Parkinson’s disease and Glaucoma. Last but not least, the time complexity of the method SSADDA is less than that of other contemporary alignment-based methods.

## 5. Conclusion

Mutations may be amino acid substitution or indel, the intensity of harmfulness may not be the same always. The effect of mutations on the stability of protein structure largely depends upon the codon positions. Determinative degree of an amino acid is considered as a passive characteristic of amino acid which shows the degree of “predeterminativity”. Determinative degree of amino acid sequence helps us to analyze new abstract amino acid “predeterminativity” properties along with known biological facts [35]. Amino acid compositions are well conserved from species to species. The genes responsible for occurring the same disease have its unique signature of amino acid occurrence frequencies in proteins. PARKIN, PINK1, DJ1 are responsible for Parkinson’s disease, whereas, mutations in MYOC and CYP1B1 cause Glaucoma. Parkinson disease (PD) is a neurodegenerative disorder and occurs due to loss of dopaminergic neurons in the nigrostriatal pathway. Whereas, Glaucoma is a progressive optic nerve degeneration, causes irreversible blindness and has been categorised within the group of neurodegenerative diseases. Although both the diseases have different causes, symptoms and genes responsible to occur are also different, but researchers found similarities of mechanistic and pathophysiologic features between Glaucoma and other diseases related to neurodegeneration like PD [30]. Reported research studies have enough evidences for glaucoma patients to have subsequent risk to get Parkinson’s disease [45]. In our present studies we have tried to build a holistic mathematical scheme to investigate the codon alteration patterns and their impact during mutation for the genes known to be responsible for any disease and applied the concept on those two diseases. The in-silico analysis states that although participating genes are separate, but the pattern of selection of codon for alteration have similarities in various ways. Hence, our scheme may help the biologists to understand the evolutionary origin of the genes participating in both the diseases. Finally, it can be concluded that our scheme gives us a more microscopic view of the existing genetic code table and thereby is able to provide information in depth when mutations take place.

## 6. Data and Software Availability

To carry out the experiment the mutations are collected from different online sources viz. http://www.molgen.vib-ua.be/PDMutDB/, http://www.myocilin.com/variants.php, https://www.omim.org/,https://www.ncbi.nlm.nih.gov/pubmed/, https://databases.lovd.nl/shared/genes/CYP1B1. The supplementary table S5 contains all the mutation specifications.

## Supporting information

*S1 Table*. The active domains that altered due to mutations.

*S2 Table*. Transition of codon groups during mutation. The highest transitions are highlighted using red colour.

*S3 Table*. Trend of alteration in density of codon with the change in codon group due to mutation (%).

*S4 Table*. Gene-wise deleterious mutations.

*S5 Table*. Mutation Specification.

## Footnotes

§ http://www.molgen.vib-ua.be/PDMutDB/

§ http://www.myocilin.com/variants.php

§ https://databases.lovd.nl/shared/genes/CYP1B1

§ https://www.omim.org/

§ https://www.ncbi.nlm.nih.gov/pubmed/

